# Distinct neural representations for prosocial and self-benefitting effort

**DOI:** 10.1101/2021.09.27.461936

**Authors:** Patricia L. Lockwood, Marco Wittmann, Hamed Nili, Mona Matsumoto-Ryan, Ayat Abdurahman, Jo Cutler, Masud Husain, Matthew A. J. Apps

**Author notes:** Correspondence should be addressed to: Patricia L. Lockwood, Centre for Human Brain Health, University of Birmingham, United Kingdom.

## Abstract

Prosocial behaviours – actions that benefit others – are central to individual and societal well-being. Most prosocial acts are effortful. Yet, how the brain encodes effort costs when actions benefit others is unknown. Here, using a combination of multivariate representational similarity analysis and model-based univariate analysis during fMRI, we reveal how the costs of prosocial efforts are processed. Strikingly, we identified a unique neural signature of effort in the anterior cingulate gyrus for prosocial acts both when choosing to help others and when exerting force for their benefit. This pattern was absent for similar self-benefitting behaviour and correlated with individual levels of empathy. In contrast, the ventral tegmental area and the ventral insula signalled subjective value preferentially when choosing whether to exert effort to benefit oneself. These findings demonstrate partially distinct brain areas guide the evaluation and exertion of effort costs when acts are prosocial or self-benefitting.

## Main

From holding open a door for a stranger to volunteering for a local charity, humans often make decisions to incur costs to benefit others^1,2^. Such ‘prosocial’ behaviours are vital for maintaining individual physical^3^ and mental health^4^ and are positively correlated with societal economic success^5^. However, whilst a plethora of research has begun to probe the psychological and neural mechanisms underlying how people make decisions about whether to donate to charity or share money, much of this work overlooks a key component: effort^6–9^. In order to behave prosocially, we have to decide whether we are willing to exert effort to help, and once committed, to energise our actions^9,10^. However, how the brain represents the effort of a prosocial act, and whether this representation is distinct from self-benefitting acts, is unknown. Uncovering such specificity is critical for connecting computational and neural levels of explanation of social behaviour^11–13^.

Effort is typically considered as costly and aversive^14–17^. If two courses of action are associated with the same rewarding outcome, most individuals will choose the course that requires less effort. This phenomenon, referred to as effort-discounting, relies on computations in which rewards are subjectively devalued by the cost of exerting effort^9,18–20^. As such, people appear only to exert effort when it is considered ‘worth it’ for the rewards that can be obtained. Research across species has begun to identify the anatomy engaged during such computations. Activity in the dorsal anterior cingulate cortex (dACC)/dorsomedial prefrontal cortex (dmPFC) and anterior insula (AI) has consistently been shown to covary with the magnitude of rewards and level of task difficulty, both prior to and during the performance of a task^18,21–25^. In addition, activity in these regions tracks subjective value during effort-based decisions^18,24–31^. Lesions to these brain areas have been linked to reductions in motivated behaviour and to a reduced willingness to exert effort^32^. Such findings implicate the dACC/dmPFC and AI as crucially engaged both when deciding whether to exert effort for reward, and when energising effortful processes.

However, existing work has typically only examined self-benefitting behaviours, where the effort is exerted to obtain rewards for one’s own benefit. Yet, the cost of effort may be different when it comes to prosocial acts. Lockwood and colleagues^9^ required participants to make decisions about whether they would rather take a rest for small reward (1 credit) or exert physical effort (30-70% of their max grip strength) to obtain higher rewards (2-10 credits). On half the trials participants chose whether to exert the effort to obtain credits for themselves, but on the other half the credits were delivered to an anonymous other person. Whilst people were willing to exert effort to obtain rewards for others, the effort cost was evaluated to be greater than when the effort was self-benefitting, and participants were less willing to exert higher levels of effort for prosocial acts^8,9,33,34^. This differential weighting of effort costs into valuations raises the possibility that partially distinct neural mechanisms may guide decisions of whether to exert effort for prosocial and self-benefitting behaviours.

Whilst there is limited research examining the neural mechanisms underlying prosocial effort, studies examining how we vicariously process others rewards or efforts implicate a potentially ‘socially’ specialised system^35,36^. Studies in which self and other trials are separated in the design, allow questions about social specificity to be addressed^12^. In such studies a sub-region of the anterior cingulate cortex lying in the gyrus (ACCg) has been implicated in processing social information. Neurophysiological recordings in monkeys indicate that the ACCg contains a higher proportion of neurons that signal exclusively when another, not oneself, receives rewards compared to other frontal cortex regions^37^. The BOLD response in this region varies as a function of the vicarious net-value of other people exerting effort, the probability and outcome of another person receiving a reward, and tracks learning about others’ ownership, but does not process similar information about one’s own effort, ownership or reward^38–41^. Activity in ACCg has also been shown to correlate with self-reported individual differences in empathy, an affective process closely linked to motivating prosocial behaviours^36,42,43^. In addition, activity in a connected portion of the temporo-parietal junction (TPJ) has long been implicated in social cognition and prosocial behaviour, and encodes effort costs differently when behaviours switch from being cooperative to competitive^44–50^. Thus a partially specialised neural circuit, comprising the ACCg and TPJ, may be engaged when deciding whether to exert effort to benefit others and applying the energy required.

Here, to address the question of whether prosocial efforts are processed distinctly from self-benefitting ones, participants completed a physical effort task^9^ where self-benefitting and prosocial decisions were dissociated. People chose between a work option and rest option on each trial whilst undergoing functional magnetic resonance imaging (fMRI). Half of the trials were self-benefitting, where they chose whether to exert effort to obtain rewards for themselves (Self), whilst the other half were prosocial - where the participant chose whether to exert effort to obtain rewards for an anonymous other person (Other, see Methods and Figure S1). If they chose to work, they needed to execute the required force to obtain that reward. Using this design, we could examine activity time-locked to the points in the trial where people made a decision to work or rest and responses during the exertion of force (Figure 1A). Participants also completed a self-report assessment of individual differences in empathy. We used a combination of parametric (and model-based) univariate analyses, as well as model-based, multivariate Representational Similarity Analysis (RSA). RSA allowed us to directly test for social and self specific representations of effort when deciding whether to benefit self and other, as well as subjective value and reward. It also enabled us to test for novel neural patterns that cannot be revealed from a purely univariate analysis, but that may be crucial for prosocial behaviour^51^.

**Figure 1.**
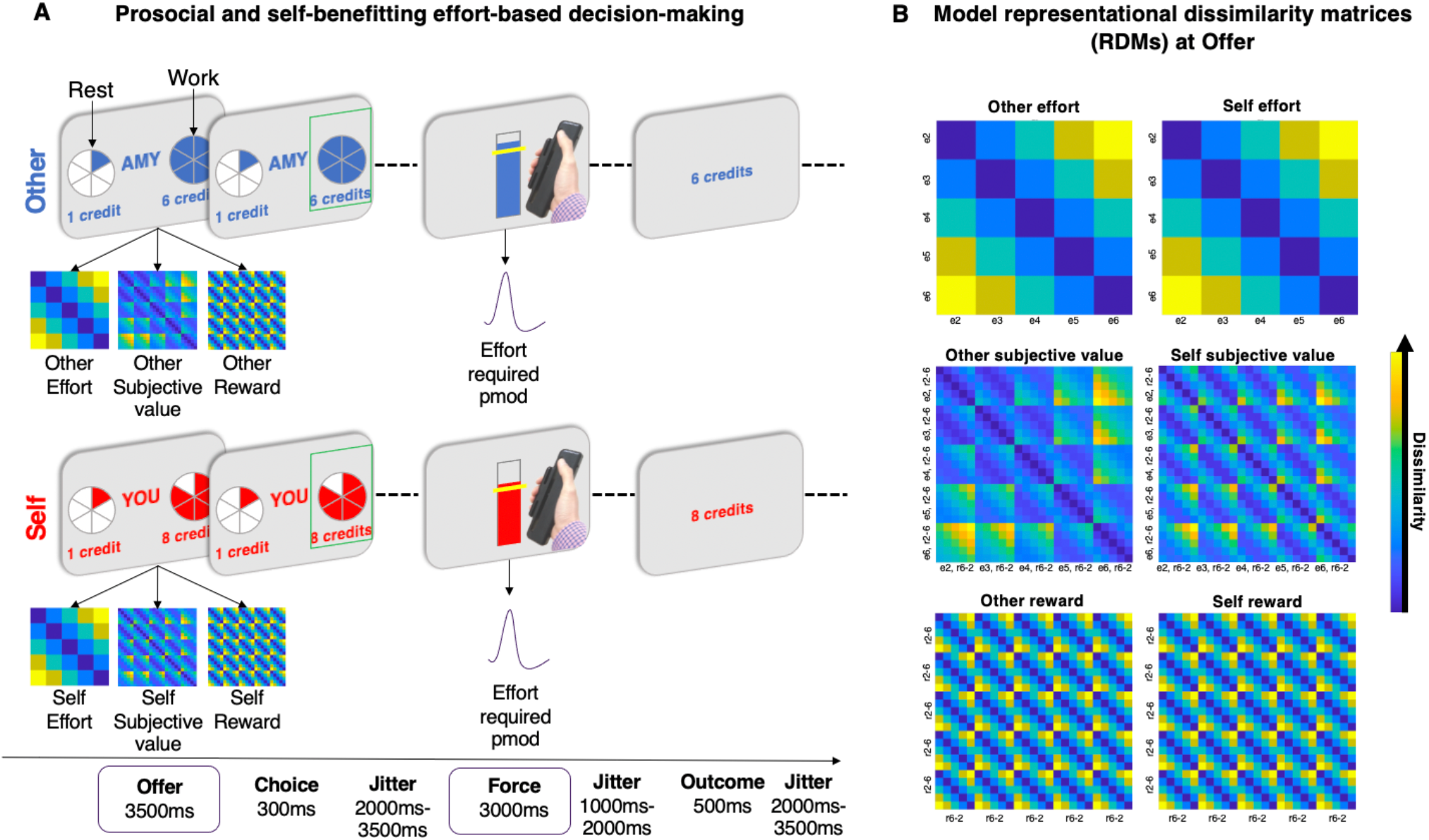
Prosocial and self-benefitting effort decision-making task and model RDMs. **(A)** Before undergoing fMRI participants were instructed to squeeze as hard as they could to measure their Maximum Voluntary Contraction (MVC) on a handheld dynamometer to threshold each effort level to their own grip strength. After thresholding and practice, participants were presented on each trial with a choice between a rest option, which required no effort (0% MVC, corresponding to one segment of the pie chart) for a low reward of 1 credit, and a work option, which required more effort (30%–70% MVC, corresponding to 2-6 segments in the pie chart) yet also generated more reward (2–10 credits). The offered reward and effort levels were orthogonal in the design. After making their selection, participants then had to exert the required force to the correct degree to receive the reward. Visual feedback of the amount of force used was displayed on the screen. Participants were informed that they would have to reach the required force level (marked by the yellow line) for at least 1s out of a 3s window. Participants then saw the outcome that corresponded to the offer they had chosen, unless they were unsuccessful, in which case “0 credits” was displayed. Crucially, on Self trials, participants made the choice, exerted the effort, and received the reward themselves, whereas on Other trials (“AMY” in this example), participants made the choice and exerted the effort, but the other participant received the reward. Participants completed 200 trials, 100 with outcomes for themselves and 100 with outcomes for the other person. Self and other trials were interleaved (see Methods). **(B)** Six 25 x 25 (5 effort and 5 reward levels) model representational dissimilarity matrices (RDMs) were constructed at the offer stage to analyse activity during effort-based decisions separately for the self and other conditions. These coded for different task features in multivariate space. For the effort model RDMs, this was the Euclidean distance between effort levels on offer. For the subjective value RDMs, this was the Euclidean distance between subjective values of offers based on the winning computational model with separate discount parameters (*Κ*) for self and other trials. Each participant’s individual *Κ* parameter was used. For the reward RDMs this was the Euclidean distance between reward levels on offer. To examine activity during the energisation of actions (force period), univariate parametric modulators (pmod, panel a) of the effort required on each trial were fitted to the onset of the force period; e: effort level, r: reward level. Yellow colours show conditions are more dissimilar whereas dark blue colours show conditions are more similar in terms of the Euclidean distance between conditions.

We show a distinct multivariate pattern of effort in the ACCg when deciding whether to act prosocially, and that activity in this region scales parametrically with the force required during exertion into prosocial but not self-benefitting acts. This pattern correlated with individual differences in self-reported affective empathy, with those highest in empathy having more distinct patterns of prosocial effort in ACCg. A domain-general set of regions in the AI and dACC/dmPFC signalled multivariate and univariate representations of subjective value for self and other. In contrast, a ventral portion of the mid-insula and the ventral tegmental area carried self-benefitting univariate and multivariate representations of subjective value, respectively. Together these results reveal partially specialised neural mechanisms for prosocial and self-benefitting efforts. These findings are important for our understanding and fostering of motivated behaviour and particularly effortful prosocial behaviours.

## Results

### People discount rewards by effort more strongly for others than for self

We analysed how peoples’ decisions to select the work offer over ‘rest’ were affected by the recipient (self or other), effort required, reward on offer, and whether participants treated prosocial decisions as distinct from self-benefitting ones (recipient). To do so we ran a generalised linear mixed-effects model (GLMM; see Methods). We observed a significant recipient*effort*reward interaction (odds ratio (OR)=0.81, 95% confidence interval=[0.69, 0.95], *p*=0.009), showing that people were less willing to help others at higher effort and lower reward levels. We also observed significant interactions between recipient and reward (OR=1.71 [1.44, 2.04], *p*=0.001), effort and reward (OR=1.23 [1.11, 1.36], *p*=0.001), and main effects of recipient, effort, and reward (Figures 2a-b, S2 and Table S1). Therefore, participants’ choices to help others were affected by the effort and reward levels on offer. In particular, participants were less willing to exert effort to reward other people than themselves.

**Figure 2.**
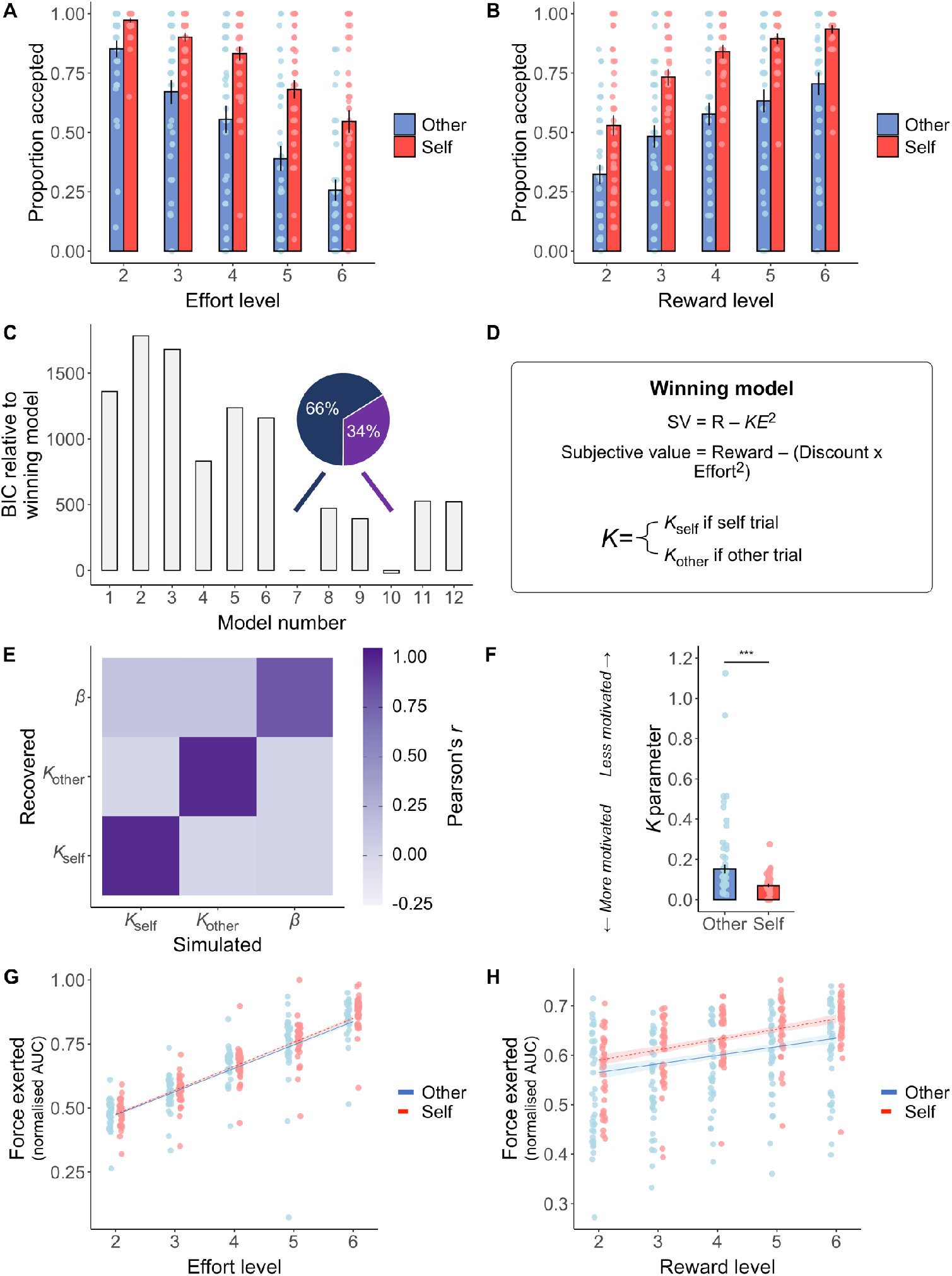
Choice, force, and computational modelling of prosocial and self-benefitting decisions. **(A)** Participants were less willing to accept the work offer over the rest offer as the effort level increased. **(B)** The proportion of work offers accepted over the baseline option increased as reward increased. Across effort and reward levels, participants were less willing to work when the other person would benefit than when they would benefit. This tendency to work more for self than others was most pronounced at the higher reward levels and particularly when a high level of effort was required. Data are represented as mean ± SE. **(C)** We compared a range of computational models of effort discounting that varied in terms of whether models had a single or separate discount (*Κ*) parameter(s) for Self and Other trials (models 1-6 vs. models 7-12) and whether the shape of the discount function was parabolic (models 1, 4, 7,10) linear (models 2, 5, 8, 11) or hyperbolic (models 3, 6, 9, 12). Models 7 and 10 had the lowest Bayesian Information Criterion (BIC) scores. These were both parabolic and had separate *Κ* parameters for self and other. However, model 7, that contained a single choice stochasticity parameter (*β*), explained behaviour in the majority of participants and was selected as the winning model. Bars show model BIC, proportions show the number of participants with the lowest BIC for model 7 compared to model 10. **(D)** Equation for the winning parabolic model with separate discount (*Κ*) parameters and a single choice stochasticity (*β*) parameter that explained behaviour in the majority of participants. **(E)** Parameter recovery using simulated data from the winning model and choice schedule showed excellent recovery of the model parameters. **(F)** Statistical comparison of the *Κ* parameters from the best fitting model showed that participants had a lower *Κ* parameter for self-benefitting compared to prosocial choices. Data are represented as median ± SE, *** shows *p*<0.001 in Wilcoxon two-sided signed rank test. **(G)** Force exerted (normalised areas under the curve during the effort period) for each of the levels of effort. Participants exerted less force for others overall and there was a significant 3-way interaction between recipient, effort and reward (*p*<.001). **(H)** Force exerted for each reward level shows participants exerted more force for higher rewards but this effect was reduced when the other person would benefit.

Next we fit and compared a range of different models of effort-discounting to participants choice behaviour using maximum likelihood estimation^9,18–20,34^. These models allowed us to test different theoretical predictions regarding the effect of effort on rewards in the task (whether discounting was linear, hyperbolic, or parabolic) and examine whether people made distinct choices on trials that benefitted themselves or the other person. The winning model in the majority of participants (66%) was a parabolic model with separate discount parameters (*Κ*_self_ and *Κ*_other_) and a single noise parameter (*β*), (see Methods, Figure 2C-F). The 2*Κ* parabolic models outperformed all linear and hyperbolic models, as well as models with only one discount parameter and had the lowest BIC scores. We also further validated our winning model in three ways. First, we calculated the median R^2^ for the model, and found the model was able to explain 92% (SD=10%) of the variance of choices. Second, we performed model identifiability analyses^52^ using simulated data and showed that our model comparison procedure was robust (see Supplementary Methods and Figure S3). Finally, parameter recovery^52,53^ tested whether the parameters from our best fitting model were recoverable in simulated data based on our schedule. We demonstrated excellent recovery of the 3 parameters (*Κ*_self_ = 98%, *Κ*_self_, 98%, *β* = 80%, Figure 2E). We therefore used parameters from the 2*Κ*1*β* model for all subsequent analyses.

We then performed statistical comparison of the estimated *Κ* parameters to compare discounting for self and other. This analysis showed that discount values for other were significantly higher for other (*Κ*_other_ median=0.15) than for self (*Κ*_self_ median=0.07, Wilcoxon two-sided signed rank test Z=−5.34, effect size *r*=0.50, [0.30, 0.65], *p*<0.001, Figure 2F). Thus, the modelling results revealed that as the required effort increases, the subjective value of decisions decreases at a higher rate when making prosocial versus self-benefitting choices.

### People exert less force when deciding to help others

A second critical aspect of helping others is that after we have decided to help, we have to energise our actions, and actually exert the effort required, to obtain the prosocial outcome. As well as being less motivated in choosing to put in effort to reward others, people may be less invigorated when helping others and exert less force into prosocial than self-benefitting actions particularly at higher effort levels9,34. To test whether people are less invigorated by prosocial acts, we used linear mixed-effects models (LMM) to predict the force that participants exerted on each trial as a function of effort, reward, and their interactions (see Methods). We observed a significant 3-way interaction between effort, reward, and recipient (*χ*^2^(16)=39.02, *p*=0.002). We also found significant interactions between recipient and reward (*χ*^2^(4)=12.49, *p*=0.022), effort and reward (*χ*^2^(16)=47.78, *p*=0.001), and main effects of recipient, effort, and reward (all *χ*^2^s>24.74, all *p*s<0.001; Figure 2G-H and Table S2). Importantly, there was no significant difference in success between self (mean=0.98) and other trials (mean=0.98, Wilcoxon two-sided signed rank test Z=−0.59, effect size *r*=0.04, [0.00, 0.26], *p*=0.55) and Bayesian evidence for no difference (BF_01_=4.35, substantial evidence in support of the null). Therefore, consistent with previous studies, participants applied less force for other benefitting than self-benefitting decisions, particularly at high effort levels, but were not less successful^9,34^.

### Prosocial and self-benefitting neural computations

To test whether there were distinct or common neural processes when making decisions to benefit self or other, we first took a multivariate, Representational Similarity Analysis (RSA) approach^51,54^. RSA can complement and add to inferences that are made based upon univariate fMRI analyses. The approach has similarities to population analyses often applied to neurophysiological recordings. As such, it can be used to link together the algorithmic and implementational levels of interpretation of fMRI data^12,55^. RSA is well suited to designs where stimuli can be processed along continuums in different dimensions, but also linked to one another via different ways^56^. As such, it was ideal for this experiment where the work offers can be parametrised in terms of effort level, reward level or combined to calculate the subjective value. This technique allowed us to examine whether patterns of responses across voxels were more or less dissimilar as a function of each of these three decision variables. Thus, we used RSA to uniquely address questions about neural representations of information that guide decision-making for oneself and others^51,57^.

We calculated brain RDMs coding for dissimilarity (correlation distance^57^) in multivariate patterns of voxels for all pairs of experimental conditions. Since the task comprised of 5 levels of effort and 5 levels reward for each recipient this resulted in a 25 x 25 matrix that was computed separately for self and other trials (Figure 1A-B, see Methods and Supplementary Methods for full details of RSA analyses). To evaluate whether these multivariate brain patterns reflected hypothesised processes, we created six behavioural model RDMs which reflected the dissimilarity in self-effort, other-effort, self-subjective value and other-subjective value (from the winning computational model), and self-reward, other-reward. Inferences were drawn by correlating each model RDM with each brain RDM using Kendall’s τA^54^. Brain RDMs were calculated using both a hypothesis driven, anatomically specific region of interest (ROI) approach (see below) and a whole-brain data-driven searchlight approach (see Methods).

In addition to the multivariate approach, we also conducted two univariate analyses. The first used the trial-by-trial subjective values from the best-fitting computational model at the time of choice, to compare whether our ROIs also carried univariate signals of subjective value. The second univariate analysis was applied to examine activity time-locked to the force period, which scaled with the amount of effort required (Figure 1A), to examine whether any brain areas distinguished between energising actions which benefit self or other. Due to the variability between participants in the number of repetitions of effort levels – which depended on participant choice behaviour – a multivariate analysis was not suitable (since the difference in number of repetitions could impose structure on the RDMs). We created a GLM including regressors time-locked to the offer cue, and to the force period, separately for self and other trials. We included parametric modulators of the required effort level time-locked to the force period, and subjective value from the winning computational model time-locked to the offer cue.

The aim of this study was to test specific hypotheses about regions that have previously been linked to guiding effort-based decisions and those linked to processing social information that could guide prosocial behaviours. Given extensive previous work on the neural systems involved in social decision-making^12,35,58^ and self-relevant effort-based decision-making^18,24–31^ we focused our fMRI analysis on four ROIs where we had strong a priori hypotheses using independent anatomical masks defined using pre-existing parcellations (see below). These masks also allowed us to probe distinct contributions of different portions of the cingulate cortex, distinguishing dorsal dACC/dmPFC (area 8m)^18,24,28^ from more ventral portions of the ACC gyrus (ACCg)^12,35,36,58^ (Figure S4, see Methods). We also included ROIs of the anterior insula (AI)^59^ and temporo-parietal junction (TPJ)^60^. In addition to these four ROIs, we conducted exploratory whole-brain analyses for completeness and considered areas significant that survived correction for multiple comparisons at the cluster level (*p*<.05, corrected for family-wise error (FWE) after thresholding at *p*<.001^61^), both in the data-driven searchlight and in the univariate analyses.

### Patterns of prosocial effort in ACCg

To test the hypothesis that there are regions that distinctly code prosocial efforts we compared multivariate patterns between self and other trials separately (Figure 1). For a region to be considered as coding prosocial, and not self-benefitting, effort, its RDM should correlate with the other-effort model RDM, and not with the self-effort model RDM, and there should be a significant difference between the strength of those correlations. Such a correlation would demonstrate that the neural patterns discriminate strongly between task conditions that vary in the levels of effort that are required to be put in for another person; but the same patterns do not vary with the differences in effort level when the decisions are about oneself. In line with our hypothesis, the ACCg ROI carried a multivariate representation of effort on prosocial trials (other-effort mean rank correlation τA ± SE: ACCg = 0.026 ± 0.009, *p*=0.005, surviving FDR correction for 24 comparisons (6 models, 4 brain areas, 2 recipients), and was the only ROI to display a significant difference between the other-effort and self-effort RDMs (Wilcoxon two-sided signed rank test Z=−2.73, effect size *r*=0.44, [0.13, 0.69], *p*=0.006, Figure 3A).

**Figure 3.**
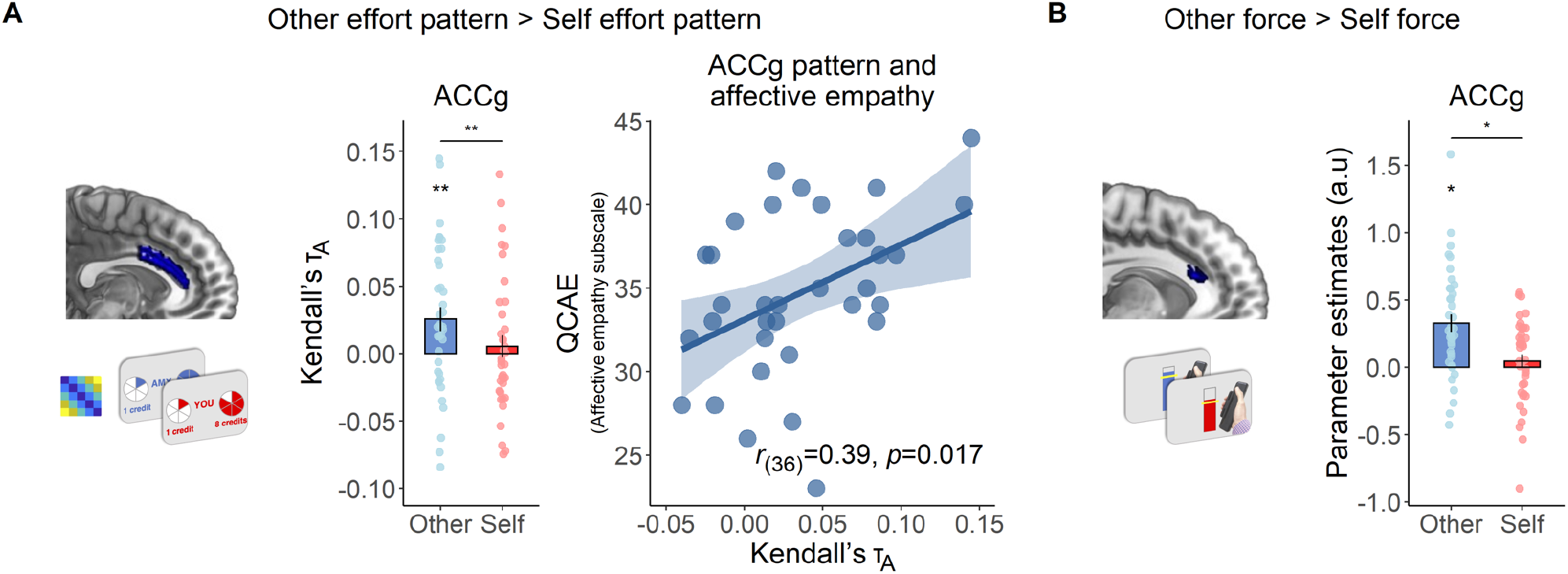
ACCg codes patterns of effort for others only, varies with level of affective empathy, and tracks effort required to benefit others only. **(A)** Across an independent structural ROI of the anterior cingulate gyrus (Neubert et al., 2014), multivoxel patterns of effort were encoded specifically for others. Kendall’s τA indicates the extent to which the effort model RDM explains pattern dissimilarity between voxels in ACCg. ACCg shows a significant correlation between the effort model RDM and the brain RDM for other (mean rank correlation τA ± SE: ACCg = 0.026 ± 0.009, *p*=0.005), as well as a greater correlation between the brain RDM and effort RDM for other compared to self (Wilcoxon two-sided signed rank test Z=− 2.73, effect size *r*=0.44, [0.13, 0.69], *p*=0.006. ROI displayed on an anatomical scan of the medial surface. Variability in ACCg effort patterns for others was explained by individual difference in affective empathy, as measured by the Questionnaire for Cognitive and Affective Empathy (QCAE^62^). In contrast, there was no significant correlation with cognitive empathy and the two corelations were significantly different from one another (*t*=2.04, *p*=0.024). (**B**) A univariate analysis time-locked to the onset of the force period showed a cluster in the ACCg that tracked the amount of effort required during the force period specifically when making prosocial but not self-benefitting decisions (x=−6, y=24, z=20, Z=3.28, k=41, *p*<0.05, FWE-SVC). Activation overlaid on an anatomical scan of the medial surface. \**p*<0.05, \*\**p*<0.01 \*\*\**p*<0.001.

Multivariate patterns in our three other ROIs also showed significant correlations with the other-effort RDM when making prosocial choices (other-effort mean rank correlation τA ± SE: TPJ = 0.033 ± 0.010, *p*=0.001; AI = 0.021 ± 0.008, *p*=0.006; dACC/dmPFC = 0.029 ± 0.008, *p*=0.001). In contrast for self-effort patterns, only the TPJ brain RDM significantly correlated with the self-effort model RDM (self-effort mean rank correlation τA ± SE: ACCg = 0.002 ± 0.009, *p*=0.61; TPJ = 0.024 ± 0.010, *p*=0.026; AI = 0.008 ± 0.009, *p*=0.40; dACC/dmPFC = 0.016 ± 0.010, *p*=0.16). Critically, although TPJ, AI and dACC/dmPFC also represented prosocial effort, they did not do so more strongly than for the self-effort RDMs (Wilcoxon two-sided signed rank test, all *p*s>0.07).

Notably the specificity for others effort in the ACCg was not due to total differences for other and self representations, as the ACCg represented other and self offers as equally dissimilar (see Methods, Bayesian paired sample t-test BF_01_=4.61, substantial evidence in support of the null). Moreover, whilst patterns in several regions significantly correlated with the other-reward RDM (other-reward mean rank correlation τA ± SE: ACCg = 0.009 ± 0.007, *p*=0.16; TPJ = 0.016 ± 0.007, *p*=0.027; AI = 0.020 ± 0.008, *p*=0.006; dACC/dmPFC = 0.025 ± 0.007, *p*=0.001), no region significantly represented others’ rewards more strongly than self-rewards (Table S3). Thus, multivariate patterns in the ACCg represented effort costs specifically when making prosocial but not self-benefitting choices (see Table S4 for exploratory whole-brain searchlight results).

### Parametric modulation of effort level when exerting force for others in ACCg

We next used univariate analysis to determine regions in which activity scaled with required effort level during the force period. We found that the BOLD response in ACCg positively covaried with force for others (x=4, y=2, z=36, Z= k=455, *p*=.001, FWE-corrected), and an overlapping cluster also showed a significant effect coding force for other > than self (x=−6, y=24, z=20, Z=3.28, k=41, *p*<.05, FWE-SVC, Figure 3B). Analysis of the force period also showed at the whole-brain level that the left TPJ positively tracked force exerted for others more than self (x=−50 y=−62 z=40, Z=4.85, k=790, *p*<.001 FWE-whole brain), with activation for other greater than self on the right side evident at small-volume corrected levels (x=52, y=− 56, z=40, Z=3.65, k=29, *p<*.05 FWE-SVC). A region in the bilateral middle insula (x=−38 y=0 z=12, Z=3.96, k=105, *p*<.05, FWE-SVC; x=44 y=4 z=10, Z=3.65, k=83, *p*<.05, FWE-SVC) tracked both self and other force exerted, but responded more strongly to other. Outside of our ROIs, we also observed significant tracking of force exerted for other more than self in a region of the superior frontal gyrus extending into the paracingulate cortex and middle temporal gyrus (Supplementary Table S5). No brain areas significantly responded more to the contrast self force>other force at the whole brain level or in any of our ROIs.

Therefore, whilst several regions processed information about prosocial efforts, only the ACCg showed a more specialised profile. Multivariate patterns in the ACCg specifically encoded representations of prosocial effort when making a choice, and univariate signals in this region scaled with how much effort was required when energising prosocial acts, with no such signals for self-benefitting efforts.

### Multivariate representations of prosocial effort in ACCg correlate with individual differences in affective empathy

Theoretical accounts suggest that empathy plays a key role in motivating prosocial behaviours^36,63,64^ with self-reported empathy a predictor of people’s willingness to exert effort to obtain rewards for others and also with responses in the ACCg when processing social information^33,36,40,65^. Thus we next sought to evaluate whether multivariate and univariate signals of prosocial effort varied with individual differences in empathy. Since we found evidence for specific effort patterns during prosocial acts in ACCg only, we focused our analysis on responses in this region. We found that affective empathy was positively correlated with the strength of correlation between the brain RDM patterns in ACCg and the other-effort RDM (Pearson’s *r*(38)=0.39, *p*=0.017, Figure 3A, right) whilst cognitive empathy was not (Pearson’s *r*(38)=0.05, *p*=0.78, correlations significantly different *t*=2.04, *p*=0.02). For the tracking of prosocial effort in ACCg during force exerted, neither affective or cognitive empathy were significantly correlated (all *r*s>-0.18, all *p*s>0.29). Thus, individuals who reported being more affectively empathic represented the effort of behaviours more distinctly in the ACCg when deciding whether to act prosocially.

### Specific coding for self-benefitting acts in midbrain and AI

Do any regions specifically code self-benefitting acts when making effort-based decisions? None of our ROIs showed a stronger correlation of the self-effort than other-effort RDM, and similarly for the SV RDM, no region showed a significantly stronger correlation for self than other (all Zs<1.37 || >1.56, all *p*s>0.12, see Table S3 for reward RDM results). However, a whole-brain exploratory searchlight analysis revealed a significantly stronger correlation with the self-SV than other-SV model RDM in the midbrain, putatively in the ventral tegmental area (VTA; x=4, y=−22, z=16, k=291, Z=4.45, *p*=.033 FWE-whole brain corrected, Figure 4A, Table S4) and in the posterior cingulate (x=20, y=−20, z=50, Z=4.78, k=578, *p*=.002, FWE-whole brain corrected, Table S4). The univariate analysis revealed a cluster in a ventral portion of the left anterior insula (vAI; x=−44 y=10 z=−10, Z=3.72, k=59, *p*<0.05 FWE-small volume) in which activity scaled more strongly with SV when making self-benefitting than other benefitting choices (Figure 4B). This cluster did not overlap with one that signalled SV on both self and other trials (see Supplementary Figure S4C). Such findings suggest that the VTA is engaged exclusively in making choices about whether it is worth exerting effort to benefit oneself, and vAI tracks subjective value more closely during self-benefitting than other-benefitting decisions.

**Figure 4.**
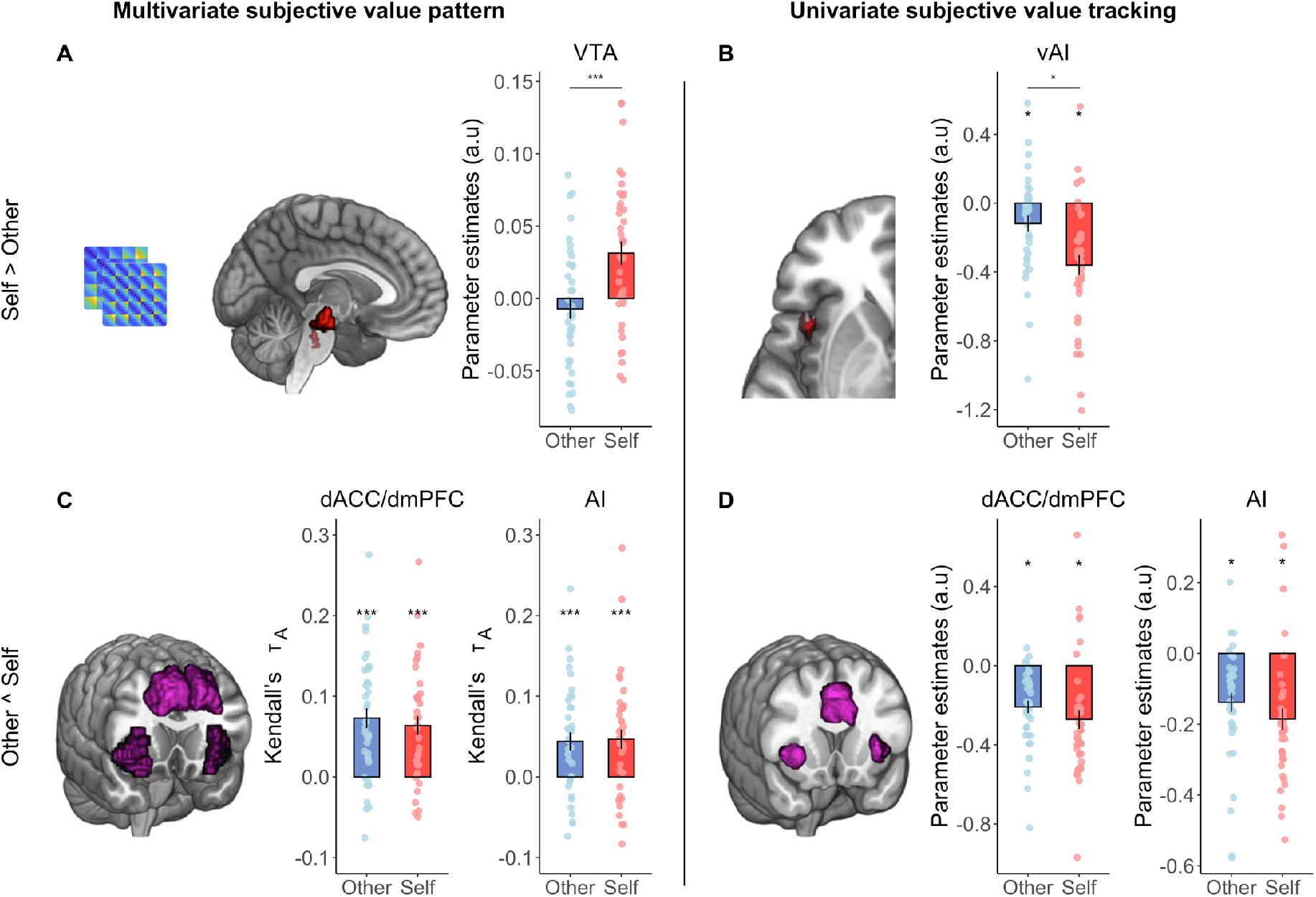
Self-benefitting and domain general representations and tracking of subjective value. **(A)** A cluster putatively in the ventral tegmental area (VTA) encoded representational patterns of subjective value exclusively on self-benefitting trials (x=4, y=−22, z=16, k=291, Z=4.45, *p*=0.033 FWE-whole brain corrected after thresholding at *p*<0.001). **(B)** A sub-region of the ventral anterior insula (vAI; x=−44 y=10 z=−10, Z=3.72, k=59, *p*<0.05 FWE-small volume) tracked subjective value of the chosen offer trial-by-trial more strongly for self-benefitting than other-benefitting choices. **(C)** The dACC/dmPFC and AI showed significant correlations between the brain RDM and subjective value RDM pattern for both other and self offers, consistent with a domain general response in these regions. **(D)** Univariate analysis also showed trial-by-trial tracking of subjective value in dACC/dmPFC (x=8 y=26 z=34, Z=4.75, k=1033, *p*<0.05 FWE-whole brain) and AI (left: x=−28 y=22 z=6, Z=4.47, k=306, *p*<0.001 FWE-SVC; right: x=34, y=24, z=2, Z=4.38, k=222, *p*<0.05 FWE-small volume) for both self and other. \**p*<0.05, \*\**p*<0.01 \*\*\**p*<0.001.

### Domain-general multivariate and univariate signals of subjective value for self and other

Previous research using univariate approaches has repeatedly implicated the dACC/dmPFC and AI in signalling SV during self-benefitting, effort-based choices in a domain-general manner regardless of the nature of the cost^18,24,28^. Next, we tested whether these regions contain information about SV when making both self-benefitting and prosocial choices (see Methods). We found significant correlations between the self-SV and other-SV RDMs in both the dACC/dmPFC and AI (self-SV mean rank correlation τA ± SE: dACC/dmPFC = 0.063 ± 0.012, *p*<0.001; AI = 0.047 ± 0.012, *p*<0.001; other-SV mean rank correlation τA ± SE: dACC/dmPFC = 0.073 ± 0.012, *p*<0.001; AI = 0.051 ± 0.013, *p*<0.001; all survive FDR correction, Figure 4C). Moreover, univariate conjunction analysis of trial-by-trial subjective value also revealed activity covarying with SV on both self and other trials in dACC/dmPFC (x=8 y=26 z=34, Z=4.75, k=1033, *p*<.05 FWE-whole brain, Figure 4D) and bilateral AI (left: x= −28 y= 22 z=6, Z=4.47, k=306, *p*<.001 FWE-SVC; right: x= 34, y=24, z=2, Z=4.38, k= 222, *p<*.05 FWE-SVC, Figure 4D) that overlapped with the same portions of dACC/dmPFC and AI that coded the multivariate pattern. Importantly neural responses to subjective value in dACC/dmPFC and AI were evident both from multivariate and univariate analyses. This is striking given that the correlation distance is invariant to the mean activation level across voxels^51,66^, rendering the multivariate and univariate predictions of neural response separate (see Methods).

Outside of dACC/dmPFC and AI, we found multivariate patterns of subjective value that overlapped between self and other in TPJ and ACCg (self-SV mean rank correlation τA ± SE: TPJ = 0.026 ± 0.011, *p*=0.018; ACCg = 0.055 ± 0.012, *p*<0.001; other-SV mean rank correlation τA ± SE: TPJ = 0.044 ± 0.011, *p*<0.001; ACCg = 0.038 ± 0.014, *p*=0.009; all survive FDR correction). At the whole-brain level, searchlight analysis also showed responses in the superior frontal gyrus, inferior parietal lobe and precentral gyrus in conjunction analyses (Table S4). No other areas significantly tracked self and other SV in any of our ROIs or at the whole-brain level. For effort, domain general patterns were observed in the bilateral precuneus (Table S4) and for reward, in bilateral precentral gyrus, cuneus, and paracentral lobule (Table S4). Univariate conjunction analysis of effort required during the force period demonstrated wide-ranging activation at the whole-brain level, centring on the precentral gyrus and cerebellum (Table S4).

## Discussion

Many prosocial acts are effortful. However, the neural mechanisms that underlie how people decide whether to exert effort into prosocial acts, and whether such mechanisms are distinct from self-benefitting acts, are poorly understood^6^. Here, we show that the ACCg processes information that is crucial for making effort-based decisions when they are prosocial, but not when they are self-benefitting. The ACCg carried a multivariate representation of effort when deciding whether to help others, and showed a univariate response to the degree of effort required whilst energising prosocial acts. The representation of effort in this area was also stronger in individuals higher in self-reported empathy. In addition, we found a region in the midbrain which processed information only when making self-benefitting effort-based choices, a portion of the ventral AI that tracked self subjective value more closely than other subjective value, as well as domain-general representations of SV in the dACC/dmPFC and distinct portions of bilateral AI. These findings highlight the importance of effort for understanding the neural mechanisms of prosocial behaviour, and that multivariate patterns and univariate model-based signals differentiate between prosocial and self-benefitting effortful acts.

There is a growing body of evidence to suggest that the ACCg is a vital cingulate sub-region for social cognition, and particularly for vicariously processing information about others^35,36^. In macaques, lesions to this region, but not neighbouring sub-regions, reduce sensitivity to social stimuli^67^. Neurophysiological recordings indicate that the ACCg contains a higher proportion of neurons that respond exclusively when seeing others obtain a reward, but not when getting rewarded oneself^37^. In rodents, a putatively homologous region contains neurons that respond when seeing another receiving electrical shocks and when exerting competitive efforts^68–70^. In humans, single-unit recordings have identified that ACCg contains neurons that signal at the time of outcomes being delivered when learning from others’, but not one’s own reward, prediction errors^71^. In addition, neuroimaging studies have shown that this region responds to cues that are predictive of rewarding outcomes for others and not self, signals the value of others’ rewarding outcomes but not one’s own when they will have to exert effort for them, and encodes social prediction errors at the time of the outcomes of others actions^35,36,41^. Combined this work suggests that the ACCg processes information about others that it does not process about ourselves. However, despite this work highlighting a role in vicariously processing others’ outcomes, whether and how this related to behaviours aimed at benefitting others, or prosocial efforts was unclear.

Crucially, we found other-specific effort effects in the ACCg that went beyond vicarious processing of others’ outcomes, and extended them to prosocial behaviours. Such findings suggest that vicarious signals in the ACCg may not be epiphenomenal or simply reflecting one’s emotional responses to others outcomes, but instead they drive behaviour^11,12^. In particular, the ACCg may be crucial for motivating people to help others, and overcome barriers to others receiving positive outcomes. Consistent with this, we found representations of effort in the ACCg both when making prosocial decisions, and when energising prosocial but not self-benefitting actions. Our finding fits with the notion that ACCg patterns do not simply reflect the reduced willingness to put in effort for others at higher effort levels, since similar self-benefitting effort representations were absent in ACCg, and those higher in empathy represented the distinction between effort costs more strongly, not less. Interpreting ACCg signals as being important for motivating actions that benefit others, rather than inhibiting them, may explain why lesions to the ACCg impair the effortful process of learning of new prosocial action-outcome associations, but not the execution of low effort previously learned prosocial acts^72,73^. Moreover, our results suggest that the finding of functional connectivity in local field potentials between the amygdala and ACCg when monkeys allocate rewards to others^74^, may be linked to ensuring the monkeys overcome any costs associated with being prosocial. Together our results indicate that the ACCg is involved in motivating the overcoming of effort costs to be prosocial, rather than being involved in effort-discounting per se.

The notion that prosocial acts can be considered as goal-directed acts, governed by similar principles and computational mechanisms as non-goal-directed actions, but implemented in distinct neural regions, highlights the need to examine effort processing to understand what makes people help others^6^. Recently, it has been shown that higher levels of affective empathy are linked to a greater willingness to exert effort to benefit others^33,65^. Here, we show that such individual variability may be linked to ACCg prosocial effort processing, with levels of affective empathy correlated with the strength of multivariate representations of prosocial effort costs when deciding whether to help others. Although the link between empathy and prosocial behaviour is often discussed, many of the mechanisms providing this link are poorly understood^36,65,75^. Our findings demonstrate that representing how costly and effortful a prosocial behaviour is, may be linked to how strongly one represents the emotional states of others, which leads to variability in how willing people are to help others.

Although we found that some regions processed information differently when making decisions about whether to exert effort to benefit self or other, several regions – and particularly the AI and dACC/dmPFC – encoded information regardless of the beneficiary. Whilst there has been some debate surrounding whether these regions encode SV in univariate fMRI studies of effort-based decision-making^18,26,30,31,76–78^, a large body of evidence suggests that lesions to these regions reduces levels of motivation^24,32^. Neurons in the dACC signal both reward value and effort costs^79^. In addition, a recent meta-analysis of fMRI studies highlights consistent evidence that these regions signal the SV during effort-based decision-making^28^. It has been argued that responses in these areas may be domain-general, with SV encoded regardless of the nature of the effort, whether it is physical or cognitive^18,28^. Our results extend this notion, as previous work has exclusively used univariate approaches. We show multivariate representations of SV are present in these same regions regardless of whom the effort is being exerted for. Such a finding supports the idea that neural processes in dACC/dmPFC and AI are an important component of motivated behaviour across multiple domains, regardless of the nature of the effort or who will receive the reward. Importantly, this also suggests that decisions to behave prosocially are distributed across both regions specialised for prosocial behaviour and those similarly involved in effort-based decisions in non-social contexts.

Notably we also found a midbrain region, closely approximating the VTA, that contained a multivariate representation of SV exclusively when making choices to benefit oneself. Previous work across species has linked the VTA to exerting effort for one’s own rewards^76,80–83^. Neurophysiological recordings in this region highlight that local field potentials are sensitive to effort requirements and that neurons in this region increase prior to deciding whether to exert effort^10,80^. Neuroimaging studies in humans suggest the VTA may be important for learning how to avoid effort costs and when deciding how much effort to allocate to a trial of a task^76,82,83^. However, our results suggest that this region may not process information for all efforts, only for those that benefit oneself. Such findings concord with the idea that prosocial and self-benefitting actions are distinct, may be linked to partially distinct motivational processes, and argue against some suggestions that a ‘warm glow’ is the primary driver of prosocial acts^8,33,34,65,75^. The warm glow account would suggest that the same mechanisms are engaged when thinking about the benefits of helping others and when benefitting oneself. Instead, we show that choosing whether to benefit ourselves may recruit some systems that are not deployed when making similar prosocial choices. This supports the idea that how prosocial one is, and one’s levels of social motivation, may be independent of how motivated one is to exert effort to benefit ourselves.

Recently we highlighted how using the framework of Marr’s three levels, can be fruitful for examining if a process is socially, or self, specific, either in how it is implemented in the brain, or in its algorithmic processes^12^. In line with this approach, our findings highlight the critical importance of breaking prosocial behaviour down into its constituent parts, for using multivariate approaches, and for designing paradigms that separate out self-benefitting from other-benefitting decision-making. Previous work examining prosocial behaviour, particularly that using economic games, has been crucial for implicating the systems that involved^1,84–86^. However, the precise computations performed have been hard to identify due to the challenge of untangling self-benefitting from other-benefitting components. We reveal that several regions, including in the AI and dACC/dmPFC which have been implicated in prosocial behaviours^48^, in fact carry information when making both self-benefitting and other-benefitting choices. Such a finding raises the possibility that these areas may be less directly linked to prosocial behaviour, and more linked to domain-general decision processes during economic games. Moreover, by measuring both self and other decision-making, we were able to identify an at least partially self-specific processes in the VTA and a different sub-region of AI, and prosocially specific representation of effort in the ACCg.

In conclusion, many prosocial acts require effort. Using computational modelling of behaviour, and multivariate and univariate analyses of fMRI data, we find evidence of distinct neural patterns of effort for prosocial and self-benefitting acts. The ACCg carries a multivariate representation of effort when making prosocial choices, and is engaged when energising prosocial acts, but does not carry similar information for self-benefitting choices or acts. In addition, we show that the AI and dACC/dmPFC all carry information that guides both self-benefitting and prosocial behaviours. Intriguingly, these areas both represent the dissimilarity between different subjective values within a neural pattern, as well as tracking the quantity of subjective value trial-by-trial. In contrast, the VTA processes the structure of subjective value only of self-benefitting acts and the ventral AI more closely tracks self-benefitting, compared to other-benefitting, values. These findings provide a unique insight into how the brain makes decisions about whether to put in effort to help others out.

## Materials and Methods

### Participants

41 healthy, right-handed participants took part. Three participants were excluded. Two who did not believe the deception in the study set-up (see “Role assignment” details below), and one who never chose to exert effort for the other person. The final group of 38 participants (26 females, mean age 23, range 18-34). Based on the effect size from Lockwood et al.,^9^ a sample of 38 people gave 83% power to detect a significant behavioural effect.

Participants were recruited through student mailing lists, online advertisements on a study recruitment board, through social media, and by word of mouth. The study was described as a social decision-making study involving pairs of participants. Participants believed that, on the day of testing, one of the pair would be randomly allocated to complete the task in the fMRI scanner whilst the other would complete the task in a testing room. In reality, all participants completed the task in the scanner, and a confederate served as the other participant. Exclusion criteria were previous study of psychology, previous participation in a study involving social interactions, left handedness, and neurological or psychiatric disorder. These questions were asked via an online screening procedure and only participants who met these criteria were invited to take part. The study was approved by the Medical Sciences Division Research Ethics Committee of the University of Oxford. Participants were paid for their participation at a rate of £15/hour, plus a bonus of up to £5 based on the credits they earned in the task. They were also told the number of credits that they earned in the prosocial condition would translate into an additional payment of up to £5 for the other participant (see details of the task below).

### Procedure

Approximately 1 week before attending the testing session, participants completed a questionnaire assessment of empathy online using the Questionnaire of Cognitive and Affective Empathy (see below for further details). Participants then attended the lab to complete a physical effort-based decision-making task modified for scanning from previous behavioural studies^9,34^. Physical effort was operationalized as the amount of force participants exerted on a handheld dynamometer. On arrival and after consent, participants were instructed to squeeze the handheld dynamometer as hard as they could. Participants were provided with visual feedback whilst doing so and encouraged to reach a line that was 110% of their maximum voluntary contraction (MVC) which they repeated for 3 trials. After this thresholding procedure, and before any task instructions, participants were introduced to another participant anonymously (see “role assignment’ procedure below). Participants practiced each of the 5 effort levels twice to ensure that they could be achieved. In the main task inside the fMRI scanner, participants were prompted to choose between one of two offers on each trial. One option allowed participants to earn a low reward for low effort (rest); the other presented a variable higher-reward, higher-effort offer (work) of the same duration. The low-reward, low-effort offer earned 1 point and required no effort. Higher-reward, higher-effort offers varied from 2-10 points (in 2-point increments). Effort ranged from 30-70% (in 10% increments) of the participants’ MVC. Participants were aware that points earned corresponded to later compensation, but were not made aware of the exchange rate while completing the task. Critically, each trial also varied in whether the outcome would be delivered the participant themselves (Self) or the receiver participant (Other, prosocial). The level of effort required for each offer was represented using coloured portions of a pie chart (Figure 1). Rewards (points) on offer for each option were written in colour below. Participants were allotted 3.5 seconds to make a choice between the rest and work offers. If they failed to choose an option, they were awarded 0 points after a full trial duration. After choosing, participants were shown a screen with a yellow horizontal bar on an empty vertical box. The horizontal bar represented the level of effort required; the box filled according to the force participants exerted on the dynamometer, providing feedback in real-time. For a trial to be considered successful, and rewards obtained, participants had to accumulate at least 1 second at or above the required force level across the 3 second force period.

The task was broken into four blocks, with a minute break in between each block to rest and prevent the build-up of fatigue. Participants selected the choice they wanted using a game controller in their left hand and used their right hand to squeeze the dynamometer. Each participant completed 100 interleaved trials per recipient (self or other).

### Role assignment

Participants were introduced to another participant who was in fact a confederate of the experimenter, as in previous studies of social decision-making^34,87^ (Figure S1). Participants were instructed not to speak and wore a glove to hide any physical characteristics and to ensure they were anonymous to one another. A second experimenter brought the confederate to the other side of the door who was also instructed not to speak and wore a glove. Participants only ever saw the gloved hand of the confederate, but they waved to each other to make it clear there was another person there (Figure S1). The experimenter tossed a coin to determine who picked a ball from the box first and then told the participants which roles they had been assigned to, based on the ball that they picked. Unbeknownst to participants, our procedure ensured that participants always ended up in the role of the person performing the effort task inside the MRI scanner and they were led to believe the other participant would be performing tasks outside of the scanner. We emphasised that the participant outside of the scanner would only perform experimental tasks that would result in outcomes for themselves, and would be unaware of the task performed by the participant inside the scanner so any reward given would be anonymous. This procedure minimised as much as possible any prosocial behaviour being due to social preferences of reciprocity^88^.

After finishing the task in the scanner, participants completed a short debriefing questionnaire where they were probed as to whether they believed they were earning rewards for another participant. Two participants reported a disbelief in the deception and were removed from analysis.

### Statistical Analysis of Behavioural Data

Analyses of behavioural data were performed using a combination of MATLAB (2019, The MathWorks Inc.) and R (version 3.6.2) using RStudio^89,90^. For choices between the work and rest offers we coded choice as a binary outcome variable and ran a generalised linear mixed-effects model (GLMM) using the *glmer* function from the lme4 package^91^. The model included fixed effects of recipient, effort (squared to mirror the computational model results), reward, and all interactions, as well as a subject-level random intercept. Squared effort and reward were z-scored before being entered into the model. We exponentiated the resulting standardised coefficients and standard error to generate odds ratios and their 95% confidence intervals. Due to the non-normal distribution of the *Κ* parameters, we compared discounting for self and other using a non-parametric Wilcoxon two-sided signed rank test and generated a standardised effect size (*r*) for this difference using the *wilcox_effsize* function from the rstatix package^92^.

For analysis of force exerted following a choice to work, we normalised participants’ force as a proportion of their maximum to account for between-subject variability in force exerted and calculated the area under the curve for the 3 second window in which they exerted force. We ran a linear mixed-effects model (LMM) that predicted normalised force with fixed effects of recipient, effort level (2, 3, 4, 5, 6), reward level (2, 3, 4, 5, 6), and all interactions, plus a subject-level random intercept. For both the GLMM on choices and LMM on force, we fit the models by maximum likelihood and tested the fixed effects for statistical significance using parametric bootstrapping (1000 simulations) with the *mixed* function from the afex package^93^. We used type II tests meaning the significance of a variable was tested by comparing the full model with the next most complex model that does not include that variable. For completeness, we also report Z statistics for the GLMM of choices and *χ*^2^ statistics for both models from these comparisons (Table S1 & S2).

### Questionnaire of Cognitive and Affective Empathy

Before the testing session, participants completed an online pre-testing questionnaire. The questionnaire aimed to measure individual levels of empathy that might influence prosocial behaviour. Empathy is the ability to vicariously experience and understand the affect of other people ^36,94^. This ability modulates people’s social behaviour and is therefore critical to social cognition and social decision-making. The Questionnaire of Cognitive and Affective Empathy (QCAE) measures two dimensions of empathy cognitive and affective^62^. Items in the QCAE corresponded to measures of cognitive empathy (such as *I can easily work out what another person might want to talk about*) or affective empathy (*I am happy when I am with a cheerful group and sad when the others are glum*). Participants rated how much each item applied to them using a 4-point Likert scale from strongly agree to strongly disagree^62^. We used Pearson correlations to test the link between QCAE scores and multivariate representations of prosocial effort in ACCg and compared correlations using the *paired.r* function from the psych package^95^.

### Computational Modelling of Behavioural Data

For modelling of choice behaviour using trial-by-trial updates, we evaluated a number of plausible models based on past research on effort discounting^18–20,34^. Models were fitted using maximum likelihood estimation using the MATLAB function fmincon^9,34^ (Supplementary Methods). For formal model comparison, we report the Bayesian information criterion (BIC) based on the log-likelihood. The winning model was a three-parameter model with separate discount (*Κ*) parameters for self and other trials and a single temperature parameter. Discount parameters (*Κ*’s) were bounded between 0 and 1.5 to ensure parameters reflected a sensible range of behaviour in the task. We conducted both parameter recovery (Figure 2E) and model identifiability to confirm the robustness of our model (see Supplementary Methods and Figure S3).

### Imaging Methods

Scanning was conducted in a Siemens Prisma 3-Tesla MRI scanner to acquire T2*-weighted echo planar imaging (EPI) volumes with a BOLD contrast BOLD. EPI volumes were acquired at a 30 degree ascending oblique angle to the AC-PC line. The angle chosen decreased the impact of susceptibility artefacts in the orbitofrontal cortex, a method validated in previous studies^96^. Acquisition parameters were as follows: voxel size 2 x 2 x 2, 1 mm gap; TE = 30 ms; repetition time = 1254ms; flip angle = 90°; field of view = 2.16 mm. A magnetization prepared rapid gradient echo (MPRAGE) sequence with 192 slices was used to obtain the structural scan (slice thickness = 1 mm; TR = 1900 ms; TE = 3.97 ms; field of view = 192 mm x 192mm; voxel size = 1 × 1 × 1 mm resolution).

### Imaging pre-processing and analyses

Data were pre-processed and analysed using SPM12 (Wellcome Department of Imaging Neuroscience, Institute of Neurology) and a standard pre-processing pipeline. Images were realigned and unwarped using a fieldmap and co-registered to the participant’s own anatomical image. The anatomical image was processed using a unified segmentation procedure combining segmentation, bias correction, and spatial normalization to the MNI template using the New Segment procedure^97^. The same normalisation parameters were then used to normalise the EPI images, which were then spatially smoothed using an isotropic Gaussian kernel at 8 mm full-width at half-maximum.

### Imaging design: multivariate analysis

Representational similarity analysis (RSA) of fMRI data was performed using SPM12 and the RSA toolbox^54^. We estimated voxel activity patterns time-locked to the offer cue for each effort, reward and recipient combination. Both the hypothesis-driven ROI analysis and searchlight analysis were based on smoothed data^98^. In addition, the GLM modelled the onset of force exertion and the onset of the outcome, the break periods, as well as 6 motion regressors. GLMs were inspected to ensure all events could be estimated independently from one another (Figure S5). For both ROI analyses and whole-brain searchlight analyses we applied multivariate noise normalisation to the voxel activity patterns to improve reliability^54,66^ and calculated the correlation distance using the pdist function in Matlab (see Supplementary materials for full RSA processing pipeline). The correlation distance metric was chosen as a measure that is magnitude insensitive to the BOLD signal, and thus makes separate predictions from the univariate trial-by-trial model-based analysis or by using alternative distance metrics such as Euclidean distance^57,66^. As in previous RSA studies^54,99–101^ the diameter of the searchlight sphere was 15 mm (approximately 100 voxels) and we used the group level mask to define the volume for the searchlight analysis. The brain searchlight maps were correlated with each behavioural RDM using Kendall’s Tau-a to parallel the ROI analysis. The resulting maps were minimally smoothed at 2mm^102^ and then inputted into three second-level flexible-factorial designs to examine effects of effort, subjective value and reward.

Formal conjunction analyses were run to determine the areas that responded across self/other recipient conditions. Comparison analyses between self and other conditions ([−1 1] for other > self or [1 −1] for self > other) were also run at each second-level design to reveal the areas that responded specifically to one condition, yielding areas of domain-specific activation. For searchlight analysis we report clusters significant at *p*<0.05, FWE-corrected for multiple comparisons, with a cluster-defining threshold of *p*<0.001, uncorrected^61^. For ROI analysis we tested whether correlations were significantly different from zero using non-parametric one-sided Wilcoxon signed-rank tests across participants and tested whether correlations for self were different from correlations for other with two-sided paired Wilcoxon signed-rank tests. All reported comparisons survived FDR correction at *p*<.05^54^. For one-sided tests this was across 24 comparisons (4 brain RDMs, 3 model RDMs, 2 recipients) and for two-sided tests between recipients, 12 comparisons (4 brain RDMs, 3 model RDMs). ROIs were constructed using anatomical masks from regions of strong a priori interest, the dmPFC/dACC^18^, anterior insula ^18^, ACCg^35,36,38,40,60^ and TPJ^45,48^.

### Imaging Design: Univariate analysis

Three event types were used to construct regressors which would be convolved with Statistical Parametric Mapping’s canonical haemodynamic response function^103^. Onsets were modelled using regressors for the choice phase, force phase, and outcome phase. Each regressor was associated with a parametric modulator. The choice phase regressor was associated with the parametric modulator of the subjective value difference of the chosen option, as in previous work^18^, force with an effort required parametric modulator (0 if no effort or if chosen the level of effort on offer), and reward with the outcome parametric modulator (the reward outcome received on each trial). Each parametric modulator was separated by recipient (self or other). The resulting design matrix had 12 columns: the first four represented self and other choices as well as their subjective values (SVs), the next four represented the self and other force exerted as well as their force (effort) parametric modulators, and the final four held self and other outcomes and their parametric modulators. Additional regressors modelled the break phase and missed trials in participants who had missed trials. First-level design matrices were inspected to ensure the different parametric modulators could be estimated with independence (Figure S6).

First-level contrast images built from the above-described design matrix focused on self and other modulators of choice SV and force. These images were then inputted into two second-level flexible-factorial designs that tested for neural regions that tracked the predicted SV during the choice period and the level of effort during the force period. Conjunction analyses were run to determine the areas that responded across self/other recipient conditions. Comparison analyses between self and other conditions (−1 1 for other > self or 1 −1 for self > other) were also run at each second-level design to reveal the areas that responded specifically to one condition, yielding areas of domain-specific activation. Analyses were reported at *p<*.05, family-wise error (FWE) corrected at the cluster level after thresholding at *p*<.001 across the whole brain or at *p<*.05 small-volume corrected at the peak voxel level, using anatomical masks from regions of strong a priori interest, the dmPFC/dACC^18^, anterior insula18, ACCg^35,36,38,40,60^ and TPJ^45,48^.

## Data and code availability

All anonymized behavioural data and code used to generate the figures can be downloaded at OSF (https://osf.io/tm45q during peer review please see https://osf.io/tm45q/?view_only=a0e9171a075549e5946b6642334e1328).

All code used to run the computational modelling can be downloaded at OSF (https://osf.io/tm45q during peer review please see https://osf.io/tm45q/?view_only=a0e9171a075549e5946b6642334e1328). Unthresholded statistical maps can be downloaded at NeuroVault (identifier to be added on publication).

## Acknowledgements

This work was supported by a Medical Research Council Fellowship (MR/P014097/1), a Christ Church Junior Research Fellowship, a Christ Church Research Centre Grant and a Jacobs Foundation Research Fellowship to P. L. Lockwood; a Biotechnology and Biological Sciences Research Council David Phillips Fellowship (BB/R010668/1) and a Wellcome Trust Institutional Strategic Support Fund grant awarded to M. A. J. Apps; and a Wellcome Trust Principal Fellowship to M. Husain; and the National Institute for Health Research Biomedical Research Centre, Oxford, United Kingdom. The Wellcome Centre for Integrative Neuroimaging is supported by core funding from the Wellcome Trust (203139/Z/16/Z).

## Author contributions

Conceptualization P.L.L. and M.A.J.A; Methodology P.L.L., M.K.W., H.N., and M.A.J.A; Investigation, P.L.L., M.M-R, and A.A.; Formal analysis; P.L.L., M.K.W., H.N., J.C., and M.A.J.A; Writing – Original Draft, P.L.L, M.M-R., and M.A.J.A; Writing – Review & Editing, P.L.L., M.K.W., H.N., M.M-R., A.A., J. C., M.H., and M.A.J.A; Funding Acquisition, P.L.L. M.A.J.A, and M. H.; Supervision, P.L.L. M.A.J.A, and M. H.

