## Supplementary material for "Distinct neural representations for prosocial and self-benefitting effort"

a

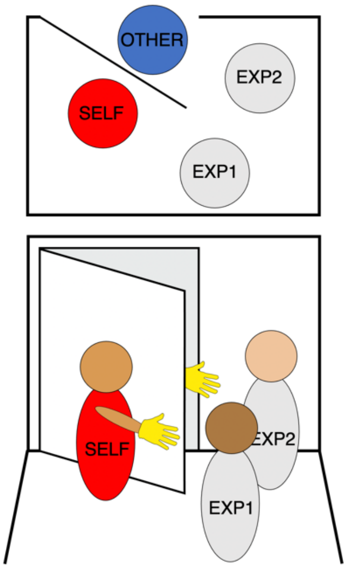


**Figure S1. Pre-task social allocation procedure. (a)** Participants were designated as “Player 1” (SELF) at the beginning of the testing session and told that they would be making decisions that impacted another player “Player 2” “OTHER” who they met at the beginning of the testing session with their identity obscured (to control for influences of identity or reciprocity, see the Method section). The procedure involved 4 people, two experimenters, EXP1 and EXP2, and two participants, “SELF” and “OTHER”.

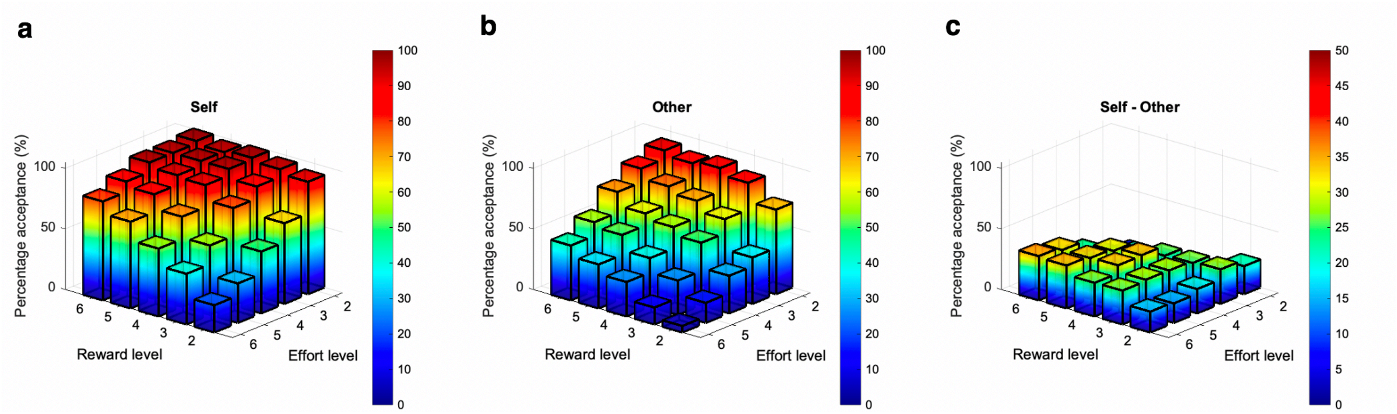


**Figure S2.** 3D plots showing percentage acceptance of work over rest offers on self and other trials and the difference between them**. (a)** Percentage acceptance on self trials as a function of reward and effort level. **(b)** Percentage acceptance on other trials as a function of reward and effort level. **(c)** Difference in percentage acceptance between self and other trials.

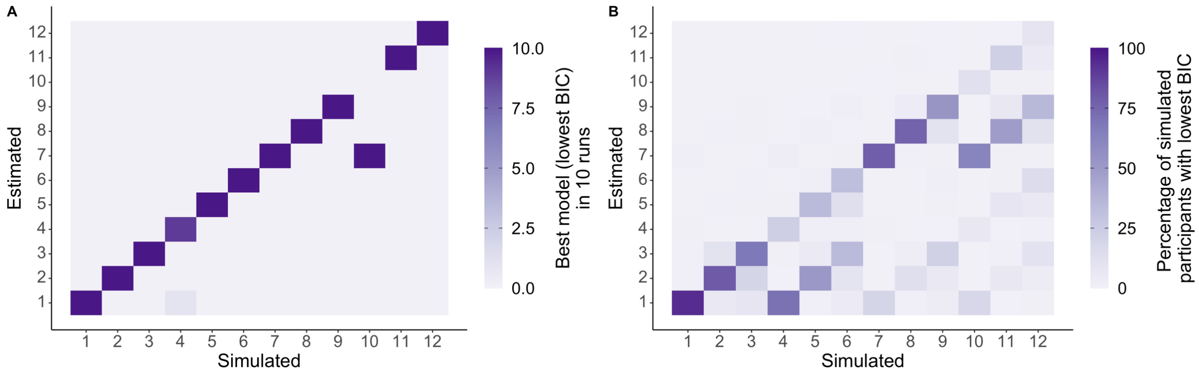


**Figure S3. Model identifiability.** Data simulated from each of the 12 models for 100 participants shows that the model comparison procedure identifies the model that simulated the data, demonstrated by the strong diagonal. **(a)** We repeated the simulations and fittings ten times and quantified the winning model as with the modelling of participants' data as the model with the lowest Bayesian Information Criterion (BIC), summing the number of times that model won across the ten runs. **(b)** We also calculated the percentage of simulated participants for which each model had the best fit to the data and averaged this over the ten runs.

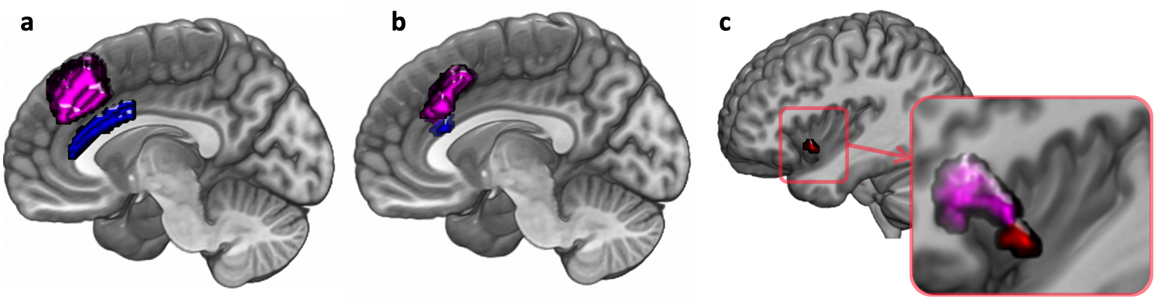


**Figure S4. (a)** anatomical regions of interest in ACCg (blue) and dACC/dmPFC (pink) overlaid on an anatomical scans of the medial surface. Notably, the two areas of the cingulate cortex are distinct. Whereas the ACCg signalled representations of effort for others only, the dACC/dmPFC signalled patterns of subjective value for both self and other. **(b)** univariate analyses show that activation in the ACCg (blue) for force exerted for others only is distinct from a separate areas of the anterior cingulate cortex in the dACC/dmPFC (pink) that negatively tracked trial-by-trial subjective value for both self and other. **(c)** univariate analysis show that activation in the ventral anterior insula (vAI) that responds more strongly on self than other trials does not overlap with the domain general portion of anterior insula (pink) that tracks subjective value in a conjunction analysis for both self and other.

**Figure S5. Correlation between regressors in the Representational Similarity analysis GLM.** A 50 x 50 GLM of regressors modelled each effort/reward combination separately for self (columns 1-25) and other (columns 26-50) trials. Within this GLM all correlations were below r<|0.007|, indicating that conditions could be appropriately estimated with independence from one another.

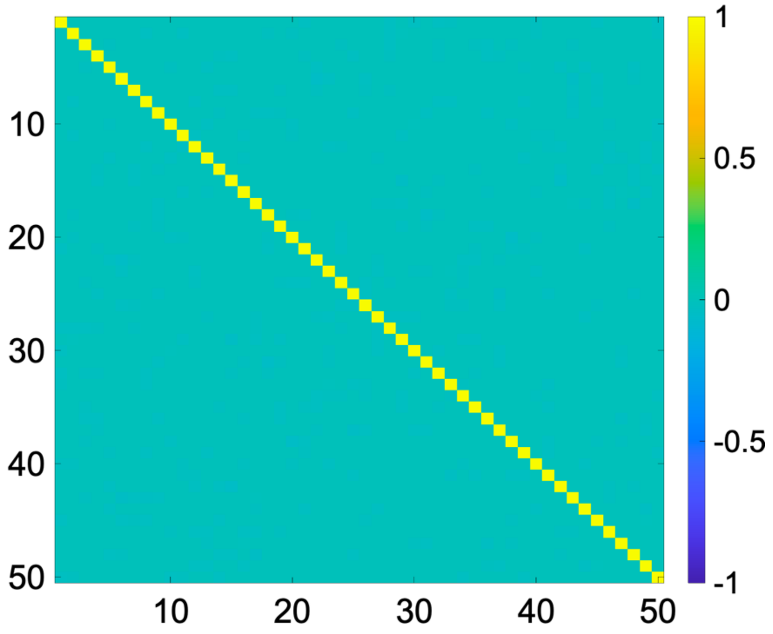

**Figure S6. Correlation between regressors in the univariate GLM**. All correlations between parametric regressors were below r<|0.178|, indicating that conditions could be appropriately estimated with independence from one another. SV = subjective value.

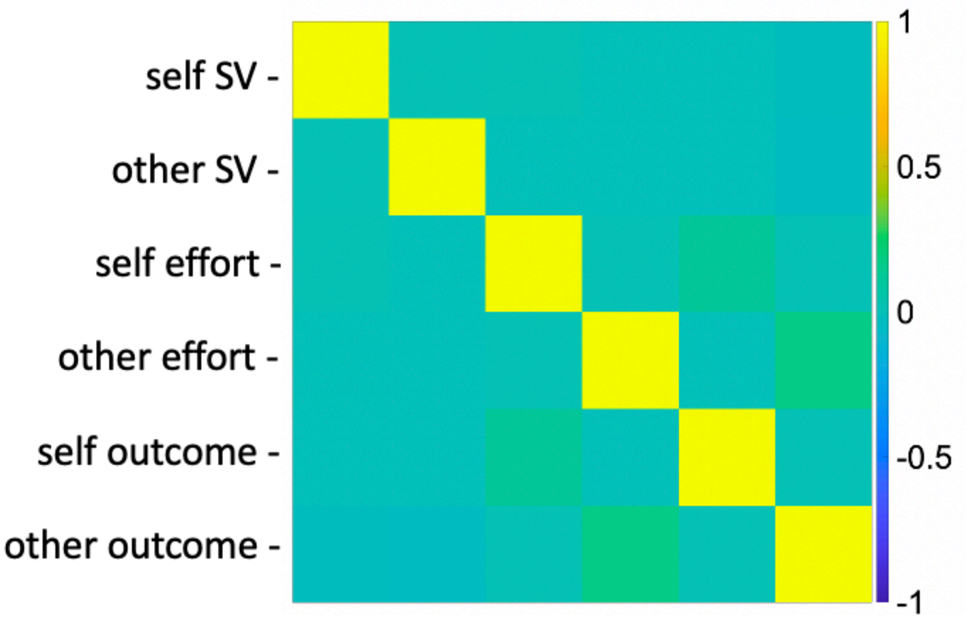

**Table S1.** Generalised linear mixed-effects model predicting choices.

| | OR | SE | CI low | CI up | Z | χ2 | df | *p* |
|---|---|---|---|---|---|---|---|---|
| (Intercept) | 1.38 | 0.36 | 0.82 | 2.32 | 1.23 | | | |
| Recipient (Self vs. Other) | 9.37 | 0.88 | 7.80 | 11.26 | 23.92 | 864.242 | 1 | 0.001 |
| Effort | 0.24 | 0.01 | 0.22 | 0.27 | -26.72 | 1702.03 | 1 | 0.001 |
| Reward | 2.54 | 0.12 | 2.31 | 2.79 | 19.29 | 1035.33 | 1 | 0.001 |
| Recipient (Self vs. Other) * Effort | 0.93 | 0.08 | 0.79 | 1.10 | -0.86 | 0.04543 | 1 | 0.85 |
| Recipient (Self vs. Other) * Reward | 1.71 | 0.15 | 1.44 | 2.04 | 6.07 | 33.6438 | 1 | 0.001 |
| Effort * Reward | 1.23 | 0.06 | 1.11 | 1.36 | 4.13 | 10.3849 | 1 | 0.001 |
| Recipient (Self vs. Other) * Effort * Reward | 0.81 | 0.07 | 0.69 | 0.95 | -2.63 | 6.95036 | 1 | 0.009 |

Note. OR: odds ratio, SE: standard error, CI: 95% confidence interval for odds ratio, low: lower CI, up: upper CI, *p* values are from type 2 tests of fixed effects using parametric bootstrapping (see Methods).

**Table S2.** Linear mixed-effects model predicting normalised force.

| | $\chi^2$ | df | *p* |
|---|---|---|---|
| Recipient (Self vs. Other) | 24.74 | 1 | 0.001 |
| Effort | 6995.26 | 4 | 0.001 |
| Reward | 66.80 | 4 | 0.001 |
| Recipient (Self vs. Other) * Effort | 7.88 | 4 | 0.11 |
| Recipient (Self vs. Other) * Reward | 12.49 | 4 | 0.022 |
| Effort * Reward | 47.78 | 16 | 0.001 |
| Recipient (Self vs. Other) * Effort * Reward | 39.02 | 16 | 0.002 |

Note. df: degrees of freedom, *p* values are from type 2 tests of fixed effects using parametric bootstrapping (see Methods).

**Table S3.** Kendall's $\tau_A$ correlations between brain RDMs and model RDMs.

| | | Mean | SE | *p* | FDR *p* |
|---|---|---|---|---|---|
| Other effort RDM | ACCg | 0.026 | 0.009 | 0.005 | 0.009 |
| | AI | 0.021 | 0.008 | 0.006 | 0.009 |
| | dACC/dmPFC | 0.029 | 0.008 | 0.001 | 0.003 |
| | TPJ | 0.033 | 0.010 | 0.001 | 0.003 |
| Self effort RDM | ACCg | 0.002 | 0.009 | 0.610 | 0.610 |
| | AI | 0.008 | 0.009 | 0.400 | 0.420 |
| | dACC/dmPFC | 0.016 | 0.012 | 0.160 | 0.170 |
| | TPJ | 0.024 | 0.013 | 0.026 | 0.032 |
| Other reward RDM | ACCg | 0.009 | 0.007 | 0.16 | 0.170 |
| | AI | 0.020 | 0.008 | 0.006 | 0.009 |
| | dACC/dmPFC | 0.025 | 0.007 | 0.001 | 0.003 |
| | TPJ | 0.016 | 0.007 | 0.027 | 0.033 |
| Self reward RDM | ACCg | 0.038 | 0.008 | <0.001 | <0.001 |
| | AI | 0.035 | 0.009 | <0.001 | 0.002 |
| | dACC/dmPFC | 0.041 | 0.009 | <0.001 | <0.001 |
| | TPJ | 0.026 | 0.009 | 0.005 | 0.008 |
| Other subjective value RDM | ACCg | 0.038 | 0.014 | 0.009 | 0.013 |
| | AI | 0.051 | 0.013 | <0.001 | <0.001 |
| | dACC/dmPFC | 0.073 | 0.011 | <0.001 | <0.001 |
| | TPJ | 0.044 | 0.012 | <0.001 | <0.001 |
| Self subjective value RDM | ACCg | 0.055 | 0.012 | <0.001 | <0.001 |
| | AI | 0.047 | 0.012 | <0.001 | <0.001 |
| | dACC/dmPFC | 0.064 | 0.012 | <0.001 | <0.001 |
| | TPJ | 0.026 | 0.011 | 0.018 | 0.024 |

Note. RDM: representational dissimilarity matrix, Mean: Kendall's $\tau_A$ correlations between brain RDMs and model RDMs, SE: standard error of the mean, FDR *p*: false-discovery rate corrected *p* value across 24 comparisons.

**Table S4**. Whole-brain RSA searchlight results.

| Brain Region | L/R | Peak voxel | | | k | t | z |
|---|---|---|---|---|---|---|---|
| **Conjunction: RSA Effort** | | | | | | | |
| Precuneus | R | 24 | -70 | 42 | 1359 | 5.55 | 5.06 |
| | R | 38 | -76 | 14 | | 4.96 | 4.60 |
| | R | 28 | -72 | 54 | | 4.26 | 4.02 |
| Precuneus | L | -18 | -74 | 50 | 409 | 4.55 | 4.26 |
| | L | -24 | -64 | 58 | | 3.92 | 3.73 |
| | L | -24 | -74 | 42 | | 3.85 | 3.66 |
| **Other > Self: RSA effort** | | | | | | | |
| *No suprathreshold voxels* | | | | | | | |
| **Self > Other: RSA effort** | | | | | | | |
| Postcentral gyrus | R | 18 | -46 | 66 | 340 | 4.84 | 4.49 |
| | R | 6 | -44 | 68 | | 4.06 | 3.85 |
| | R | 12 | -58 | 64 | | 3.72 | 3.55 |
| **Conjunction: RSA Subjective value** | | | | | | | |
| Precentral gyrus | L | -38 | -22 | 56 | 5941 | 4.85 | 4.50 |
| *ext. suppl. motor area* | L | -10 | -6 | 62 | | 4.83 | 4.49 |
| | L | -10 | 6 | 58 | | 4.80 | 4.46 |
| Superior frontal gyrus | L | -22 | 36 | 32 | 731 | 4.80 | 4.46 |
| | L | -22 | 46 | 30 | | 4.29 | 4.04 |
| *ext. dorsomedial prefrontal cortex* | L | -20 | 40 | 24 | | 4.27 | 4.03 |
| Anterior insula | L | -46 | 18 | 6 | 563 | 4.11 | 3.89 |
| *ext. Inferior frontal gyrus* | L | -56 | 16 | 22 | | 4.10 | 3.88 |
| | L | -32 | 30 | 10 | | 3.99 | 3.79 |
| Inferior parietal lobe | L | -48 | -46 | 50 | 278 | 4.02 | 3.81 |
| *ext. superior parietal lobe* | L | -32 | -56 | 56 | | 3.93 | 3.74 |
| | L | -40 | -58 | 56 | | 3.79 | 3.61 |
| **Other > Self: RSA subjective value** | | | | | | | |
| *No suprathreshold voxels* | | | | | | | |
| **Self> Other: RSA subjective value** | | | | | | | |
| Posterior cingulate | R | 20 | -20 | 50 | 578 | 4.78 | 4.45 |
| | R | 22 | -24 | 42 | | 4.76 | 4.43 |
| *ext. Posterior insula* | R | 32 | -24 | 24 | | 4.41 | 4.14 |
| Midbrain/ventral tegmental area | R | 4 | -22 | -16 | 291 | 4.44 | 4.16 |
| | L | -2 | -28 | -18 | | 4.28 | 4.04 |
| | L | -10 | -30 | -28 | | 3.75 | 3.57 |
| **Conjunction: RSA Reward** | | | | | | | |
| Precentral gyrus | R | 30 | -14 | 56 | 1058 | 5.40 | 4.94 |
| | R | 30 | -22 | 58 | | 4.90 | 4.55 |
| | R | 34 | -20 | 46 | | 4.58 | 4.29 |
| Cuneus | L | -12 | -76 | -22 | 2281 | 5.32 | 4.88 |
| *ext. lingual gyrus* | L | -20 | -82 | -14 | | 5.04 | 4.66 |
| | L | -10 | -86 | -14 | | 4.67 | 4.36 |
| Precentral gyrus | L | -42 | -14 | 60 | 1991 | 4.95 | 4.58 |
| *ext. inferior frontal gyrus* | L | -38 | 18 | 30 | | 4.42 | 4.15 |
| *ext. middle frontal gyrus* | L | -38 | 4 | 52 | | 4.39 | 4.12 |
| Superior parietal lobe | L | -22 | -66 | 58 | 319 | 4.45 | 4.18 |
| *ext. Inferior parietal lobe* | L | -28 | -52 | 48 | | 3.61 | 3.46 |
| Paracentral lobule | R | 2 | -42 | 58 | 285 | 4.30 | 4.05 |
| *ext. Precuneus* | L | -6 | -52 | 56 | | 3.59 | 3.44 |
| | L | -6 | -38 | 52 | | 3.52 | 3.38 |
| **Other > Self: RSA reward** | | | | | | | |
| *No suprathreshold voxels* | | | | | | | |
| **Self > Other: RSA effort** | | | | | | | |
| Precuneus | R | 34 | -78 | -40 | 312 | 4.59 | 4.29 |
| | R | 16 | -68 | -48 | | 4.54 | 4.25 |
| | R | 26 | -76 | -42 | | 4.32 | 4.07 |

Whole-brain searchlight results. For all regions, FWE $P < 0.05$ cluster-level whole-brain corrected after thresholding at $p<.001$. ext, extending into; k, cluster extent; L, left; PE, R, right.

**Table S5.** Whole-brain univariate results.

| Brain Region | L/R | Peak voxel | | | k | t | z |
|---|---|---|---|---|---|---|---|
| **Conjunction: univariate subjective value** | | | | | | | |
| Dorsal anterior cingulate cortex | R | 8 | 26 | 34 | 1359 | 5.16 | 4.75 |
| | L | -6 | 16 | 46 | | 5.14 | 4.74 |
| | L | -6 | 28 | 34 | | 4.70 | 4.38 |
| **Other > Self: univariate subjective value** | | | | | | | |
| *No suprathreshold voxels* | | | | | | | |
| **Self > Other: univariate subjective value** | | | | | | | |
| *No suprathreshold voxels* | | | | | | | |
| **Conjunction: univariate force required** | | | | | | | |
| Precentral gyrus | L | -20 | -28 | 64 | 116518 | 15.63 | >8 |
| *ext. cerebellum* | L | 2 | -70 | -38 | | 14.10 | >8 |
| | L | 2 | -60 | -12 | | 13.88 | >8 |
| **Other > Self: univariate force required** | | | | | | | |
| Posterior parietal lobe (temporo-parietal junction) | L | -50 | -62 | 40 | 776 | 5.28 | 4.85 |
| | L | -26 | -68 | 56 | | 3.57 | 3.42 |
| Superior frontal gyrus (dorsomedial prefrotnal cortex) | L | -12 | 26 | 52 | 2133 | 4.91 | 4.55 |
| | R | 12 | 48 | 38 | | 4.86 | 4.51 |
| | L | -16 | 50 | 28 | | 4.51 | 4.22 |
| Superior temporal gyrus | L | -56 | -38 | -8 | 414 | 4.51 | 4.22 |
| | L | -52 | -46 | -4 | | 4.32 | 4.07 |
| | L | -44 | -34 | -6 | | 4.18 | 3.95 |

Whole-brain univariate results for parametric modulators of subjective value (at offer) and force required (at onset of effort). For all regions, FWE P < 0.05 cluster-level whole-brain corrected after thresholding at p<.001. ext, extending into; k, cluster extent; L, left; PE, R, right.

### Supplementary methods

#### Computational modelling

We compared a range of models based on those validated in previous studies of effort and reward discounting[1–5]. The model space tested varied the shape of the discount function of subjective value, of choosing the more effortful option over the rest option (either parabolic (models 1,4,7,10), linear (models 2,5,8,11), and hyperbolic (models 3,6,9,12). We also compared models with single and separate noise ($\beta$) parameters and whether the same or a different discount parameter was needed for self and other (models 1-6 vs. 7-12). This resulted in 12 putative models (3 different discount functions, separate or the same discount parameters for self and other, and separate or the same noise parameters for self and other). The winning model (model 7) that explained behaviour in the majority of participants was a parabolic model with separate discount ($K$) parameters and a single noise ($\beta$) parameter. This model was very close in BIC value to another model (model 10) also with separate $K$ parameters, but separate noise parameters (Model 7 $2K1\beta$ BIC = 4,7948 vs Model 10 BIC = 4,7732 $2K2\beta$). However, the $2K2\beta$ model only won in 33% of participants. We therefore selected model 7 as the winning model, and further showed that model 7 had parameters that were recoverable and identifiable (see below).

#### Parameter recovery

Parameter recovery was performed on data simulated by the winning $2K1\beta$ model from 22,500 synthetic participants. We used a wide range of parameter values from a grid of values in the ranges: $K_{self}$ = [0:0.1:1.5]; $K_{other}$ = [0:0.1:1.5]; $\beta$ = [0:0.1:10], creating 25,856 combinations. We added noise to each of the three parameters for each simulated agent (from a standard normal distribution multiplied by 0.05) to improve our coverage of possible parameter values. After generating the simulated behaviour, we refitted the simulated behaviour using fmincon in MATLAB (2019, The MathWorks Inc.). We used the best fit from 10 random starting configurations to avoid local minima. The correlations between the true simulated and fitted parameter values were: $K_{self}$ = 0.98; $K_{other}$ = 0.98; $\beta$=0.80. Thus, parameter recovery was reliable for all parameters.

#### Model identifiability

Data were simulated from 100 synthetic participants with each of our 12 models with parameter values drawn randomly from the ranges used for parameter recovery. For example, simulated data from the $2K1\beta$ used two randomly generated $K$ parameters and one $\beta$ parameter. We then fit the models to these data in the same way as the participant data and repeated the simulation and fitting process ten times for all 12 models. On each of the ten rounds, we designated the winning model as the one with the lowest total BIC across participants and also calculated the percentage of participants for which each model had the lowest BIC. Strong model identifiability is show by the model that simulated the data winning most often (summed across the ten rounds; Figure S3a) and being best for a large percentage of participants (averaged across the ten rounds; Figure S3b).

**Representational similarity analysis**

RSA analyses were conducted using a-priori regions of interest using independent anatomical masks comprising ACCg, TPJ, dACC/dmPFC and AI[6,7]. We also conducted an exploratory whole-brain searchlight analysis. For both analyses, ROI and whole-brain RDMs were calculated using the correlation distance between all conditions[8]. For the ROI analysis, anatomical masks were realigned to be in the same voxel space as participant scans and then custom scripts were used to calculate the resulting representational dissimilarity matrices in a particular ROI with regression coefficients were spatially pre-whitened.

For the searchlight analysis[9], the searchlight was constructed around each cortical voxel, including the 100 cortical voxels with the smallest surface-wise geodesic distance from the central voxel. The searchlight was also performed using adapted scripts from the RSA toolbox ([10] original surface-based searchlight scripts from by Joern Diedrichsen and Naveed Ejaz, code available at https://github.com/rsagroup/rsatoolbox). The searchlight definition was conducted using Freesurfer's reconall command and depended on cortical reconstruction and alignment[11–13]. This procedure incorporated subject-specific anatomy by defining cortical searchlights on the 2D surface. As in the ROI analysis, regression coefficients were spatially pre-whitened within the searchlight using the RSA toolbox. These brain RDMs were computed separately for other and self trials so that they could be statistically compared. We then took each resulting 25 x 25 brain RDM and correlated it voxel by voxel with each model RDM to produce a statistical map across the whole brain, which was then saved as an image for subsequent statistical analysis in SPM. This procedure resulted in 6 brain-model RDM maps (other effort, self effort, other subjective value, self subjective value) which were then smoothed with 2mm (to allow for statistical inference within the context of random field theory[14] and input into second-level flexible factorial designs to test for the formal conjunction between other and self and any self-other differences. As in the univariate analysis, results were thresholded at p<.001 and considered significant at p<.05 whole-brain corrected at the cluster level[15].

We also conducted a control analysis to examine whether representations on self and other trials were overall more similar or dissimilar from one another in our ROI brain RDMs. For this analysis we averaged every cell in the lower triangle of the 25 x 25 brain RDM for self and other and compared the resulting estimate of dissimilarity. We found that self and other representations were equally dissimilar in ACCg ($BF_{01}$=4.21, substantial evidence in support of the null), but significantly more dissimilar for self compared to other in TPJ ($t_{(37)}$=5.07, $p$<.001), dACC/dmPFC ($t_{(37)}$=4.32, $p$<.001) and AI ($t_{(37)}$=2.79, $p$=.008).
